# Modular origins of high-amplitude co-fluctuations in fine-scale functional connectivity dynamics

**DOI:** 10.1101/2021.05.16.444357

**Authors:** Maria Pope, Makoto Fukushima, Richard F. Betzel, Olaf Sporns

**Affiliations:** Program in Neuroscience, Indiana University, Bloomington, USA; School of Informatics, Computing and Engineering, Indiana University, Bloomington, USA; Division of Information Science, Graduate School of Science and Technology, Nara Institute of Science and Technology, Ikoma, Nara, Japan; Data Science Center, Nara Institute of Science and Technology, Ikoma, Nara, Japan; Center for Information and Neural Networks, National Institute of Information and Communications Technology, Suita, Osaka, Japan; Department of Psychological and Brain Sciences, Indiana University, Bloomington, USA; Cognitive Science Program, Indiana University, Bloomington, USA; Network Science Institute, Indiana University, Bloomington, USA

**Keywords:** Connectomics, Resting State, fMRI, Brain Dynamics, Computational Neuroscience

## Abstract

The topology of structural brain networks shapes brain dynamics, including the correlation structure of brain activity (functional connectivity) as estimated from functional neuroimaging data. Empirical studies have shown that functional connectivity fluctuates over time, exhibiting patterns that vary in the spatial arrangement of correlations among segregated functional systems. Recently, an exact decomposition of functional connectivity into frame-wise contributions has revealed fine-scale dynamics that are punctuated by brief and intermittent episodes (events) of high-amplitude co-fluctuations involving large sets of brain regions. Their origin is currently unclear. Here, we demonstrate that similar episodes readily appear *in silico* using computational simulations of whole-brain dynamics. As in empirical data, simulated events contribute disproportionately to long-time functional connectivity, involve recurrence of patterned co-fluctuations, and can be clustered into distinct families. Importantly, comparison of event-related patterns of co-fluctuations to underlying patterns of structural connectivity reveals that modular organization present in the coupling matrix shape patterns of event-related co-fluctuations. Our work suggests that brief, intermittent events in functional dynamics are partly shaped by modular organization of structural connectivity.

**Significance Statement:** Brain regions engage in complex patterns of activation and co-activation over time. Relating these patterns to rest or task-related neural processing is a central challenge in cognitive neuroscience. Recent work has identified brief intermittent bursts of brain-wide signal co-fluctuations, called events, and shown that events drive functional connectivity. The origins of events are still unclear. Here, we address this gap in knowledge by implementing computational models of neural oscillators coupled by anatomical connections derived from maps of the human cerebral cortex. Analysis of the emerging large-scale brain dynamics reveals brief episodes with high system-wide signal amplitudes. Simulated events closely correspond to those seen recently in empirical recordings. Notably, simulated events are significantly aligned with underlying structural modules, thus suggesting an important role of modular network organization.

## Introduction

Structural and functional brain networks exhibit complex topology and temporal dynamics [1–3]. The topological organization of structural connectivity (SC; the connectome) is characterized by broad degree distributions, hubs linked into cores and rich clubs, and multi-scale modularity [4–6]. Functional connectivity (FC), as measured with resting-state functional magnetic resonance imaging (fMRI), displays consistent system-level architecture [7–9], as well as fluctuating dynamics [10–12] and complex spatiotemporal state transitions [13,14]. Resting brain dynamics exhibit metastable behavior, with noise-driven explorations of a large repertoire of network states and configurations [15].

Recent work has uncovered fine-scale dynamics of functional connectivity as measured with fMRI during rest and passive movie-watching [16,17]. The approach leverages an exact decomposition of averaged FC estimates into patterns of edge co-fluctuations resolved at the time scale of single image frames [18]. These studies reveal that ongoing activity is punctuated by brief, intermittent, high-amplitude bursts of brain-wide co-fluctuations of the blood oxygenation level dependent (BOLD) signal. These episodes, referred to as “events”, drive long-time estimates of FC and represent patterns with consistent topography across time and across individuals [16,17,19]. The occurrence of events appears unrelated to non-neuronal physiological processes, head motion, or acquisition artifacts. A better understanding of how events originate may illuminate the basis for individual differences in FC and its variation across cognitive state, development, and disorders. Here, we aim to provide a generative model for the origin of events in neuronal time series and uncover potential structural bases for their emergence in fine-scale dynamics.

The relationship of structure to function has been a central objective of numerous empirical and computational studies, leveraging cellular population recordings [20,21], electrophysiological [22] and neuroimaging techniques [23–25]. While there is broad consensus that ‘structure shapes function’ on long time scales [26,27] relating specific dynamic features to the topology of the underlying structural network is an open problem. Computational models have made important contributions to understanding how structural connectivity [28,29], time delays and noisy fluctuations [30] contribute to patterns of functional connectivity as estimated over long- and short-time scales. Model implementations range from biophysically based neural mass models to much simpler phase oscillators such as the Kuramoto model [31]. Despite their overt simplicity, phase oscillator models can generate a wide range of complex synchronization and coordination states, and they reproduce patterns of empirical FC [32], including temporal dynamics at intermediate time scales [33]. These modeled dynamics reproduce ongoing fluctuations between integrated (less modular) and segregated (more modular) network states [34,35], a key characteristic of empirical fMRI resting-state dynamics [36].

Here, we pursue a computational modeling approach that seeks to relate high amplitude co-fluctuations to whole-brain network structure. We simulate spontaneous BOLD signal dynamics on an empirical structural connectivity matrix of the human cerebral cortex using an implementation of a coupled phase oscillator model incorporating phase-delays, the Kuramoto-Sakaguchi (KS) model [37]. The KS model is well-suited for this purpose because its parsimonious parametrization allows for drawing specific links between network structure and synchronization patterns. We find that over broad parameter ranges BOLD signals exhibits significant high-amplitude network-wide fluctuations strongly resembling intermittent events observed in empirical data. Model dynamics reproduce several key characteristics of empirical events, including their strong contribution to long-time averages of FC as well as recurrent patterns across time. Simulated events are significantly related to network structure. They fall into distinct clusters aligned with different combinations of modules in underlying structural connectivity. Disruption of structural modules largely abolishes the occurrence of events in BOLD dynamics. These findings suggest a modular origin of high-amplitude co-fluctuations in fine-scale functional connectivity dynamics.

## Results

Empirical data were derived from 95 subjects included as part of the Human Connectome Project (HCP), with SC reconstructed from diffusion imaging and tractography and FC derived from 4 resting-state scans (792 seconds, 1100 frames, TR = 720 ms). The brain was parcellated into 200 functionally defined regions of approximately equal size [38], covering both cerebral hemispheres but excluding subcortical or cerebellar structures. SC data comprised estimates of connection weights as well as tract lengths used to compute time delays based on a uniform conduction velocity. Most simulations employed a consensus average of SC across all 95 subjects that preserved mean density as well as the distance-distribution of tracts [39]. Ten structural modules (labeled M1, M2, … M10; SI Fig 1) were derived from an implementation of multi-resolution community clustering (see Methods).

The computational model implemented the Kuramoto-Sakaguchi phase-delay oscillator ([37]; Fig 1A,B), simulated at 1 ms time resolution. A frustration matrix derived from empirical connection lengths carried phase delays computed from connection lengths and a biophysically realistic conduction velocity. Oscillator signal amplitudes (Fig. 1C) were convolved with a standard haemodynamic response function (HRF; Fig. 1D) to yield simulated BOLD signals, which were then lowpass filtered (cutoff at 0.25 Hz) and down-sampled to match the fMRI sampling rate at the empirical TR used in the HCP data. For a single simulation run, the resulting time series extended over 792 seconds, composed of 1100 time steps (720 ms). An initial transient of 20 seconds was discarded. Simulated BOLD time courses were processed into edge time series, computed as the element-wise product of the edge’s BOLD time series standard scores (Fig. 1E,F). The mean across all time steps of a single edge in the edge time series corresponded exactly to that edge’s functional connectivity [16–19]. The ‘root sum square’ (RSS) of edgewise co-fluctuations, computed across all edges on each time step, recorded system-wide instantaneous co-fluctuations (Fig. 1G). Events were detected by applying a non-parametric permutation-based null model that randomly offset time series relative to each other (Fig 1G), thus preserving (approximately) each time series’ autocorrelation while randomizing their temporal alignment (cross-correlation).

**Figure 1:**
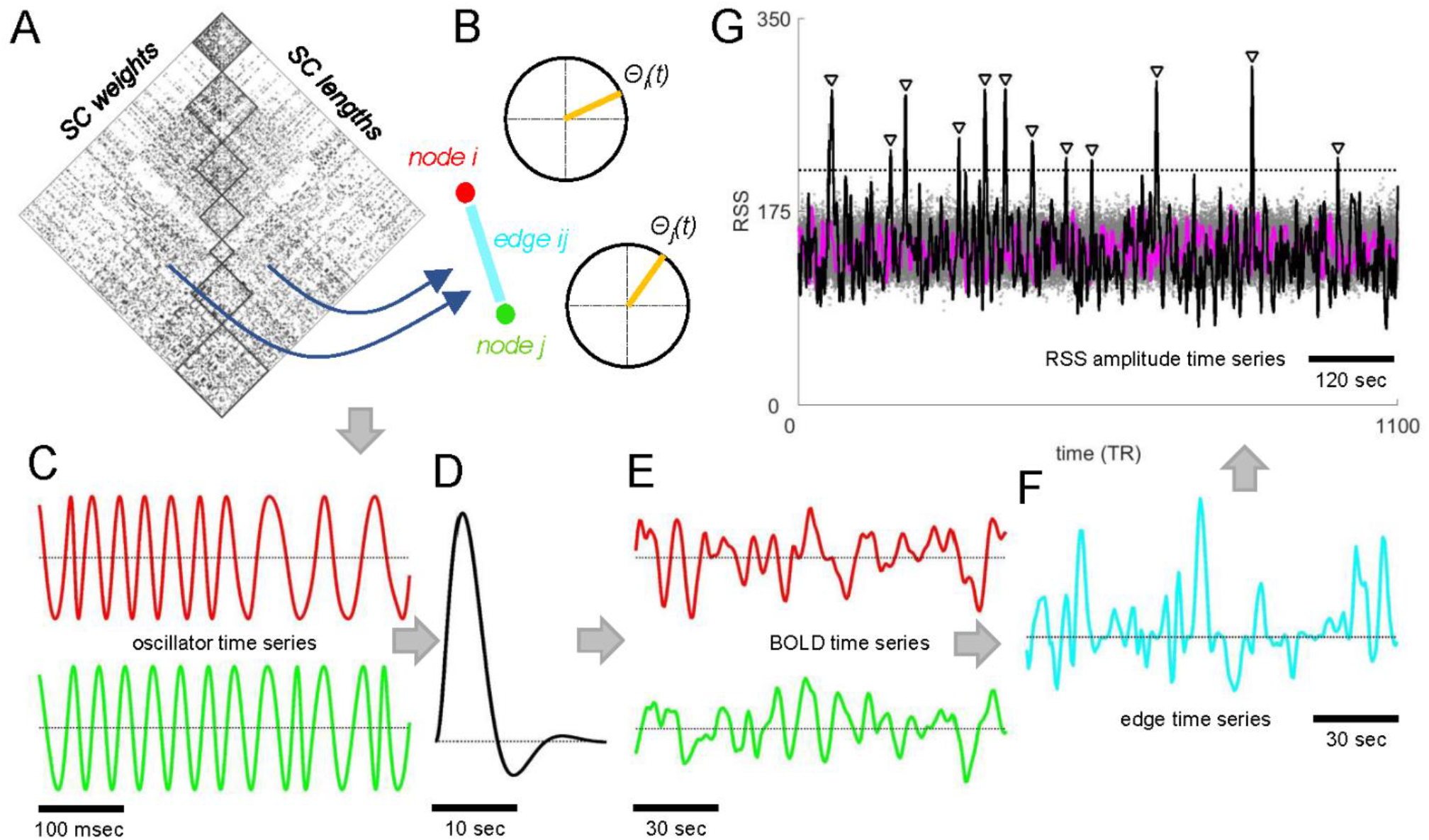
Kuramoto-Sakaguchi (KS) model schematic, computational workflow, and event detection. (A) SC weight and length matrix. (B) Node pair *(i,j)* linked by an edge *ij* and its corresponding phases *θ*(*t*). (C) Oscillator time series (*θ*(*t*)). (D) Haemodynamic response function (HRF) used for convolving oscillator time series to yield BOLD time series (E). (F) Elementwise product of normalized BOLD time series yields edge time series. (G) Root-sum-square (RSS) of all edge time series yields RSS amplitude time series. The null model consists of a distribution of null RSS amplitudes computed from randomly shifted node time series. Gray dots show amplitudes from 100 null models, stippled line indicates the *p* <0.001 cutoff derived from 1000 permutation nulls. Peaks exceeding the cut-off indicated by inverted triangles correspond to events. Data shown here computed from a representative run (*k*=280, 12 events detected).

Most simulations were carried out using a group-averaged SC coupling matrix (Fig 2A) while varying the global coupling parameter *k*, yielding simulated BOLD time series and average FC patterns. Comparison of empirical (Fig 2A) to simulated FC (Fig 2B) exhibited modest levels of similarity (Fig 2B,C), in line with previous studies. A single value of the coupling parameter (*k*=280), which provided a near-optimal match with empirical FC, was selected for further detailed analysis. Empirical FC was significantly correlated with the strengths of the SC weights (Pearson correlation, all node pairs: *r*_*a*_=0.214, *p* = 0; Pearson correlation, intra-hemispheric node pairs: *r*_*i*_ =0.343, *p* = 0; Fig 2C), as well as the co-classification matrix summarizing multi-scale modular organization of SC (Fig 2B). The latter finding agrees with prior observations in modeling studies that have linked structural to functional modules [28,32] and expands on these observations by demonstrating that SC multi-scale co-classification alone can significantly predict the long-time covariance structure of FC derived from empirical (*r*_*a*_=0.257, *p* = 0, *r*_*i*_ = 0.359, *p* =0) and modeled time series (*r*_*a*_=0.756, *p* =0 *r*_*i*_ =0.732, *p* = 0; *k* = 280). Model fit remained significant when simulations were carried out on SC matrices of individual participants (*r*_*a*_=0.200, *p* = 0 *r*_*i*_ =0.341, *p* =0; 10 randomly selected subjects, 4 simulations each; correlations computed against the mean of the same subject’s empirical FC). Realistic FC patterns emerged with only partial synchronization of phase oscillators, as documented by relatively low means of the order parameter (*mean*(*R*(*t*)) = 0.0825 ± 0.0002; SD; 12 runs, *k*=280), accompanied by consistently high standard deviations (*std*(*R*(*t*)) = 0.0825 ± 0.0002; SD; 12 runs, *k*=280), indicative of persistent variability in synchronization patterns and metastability.

**Figure 2:**
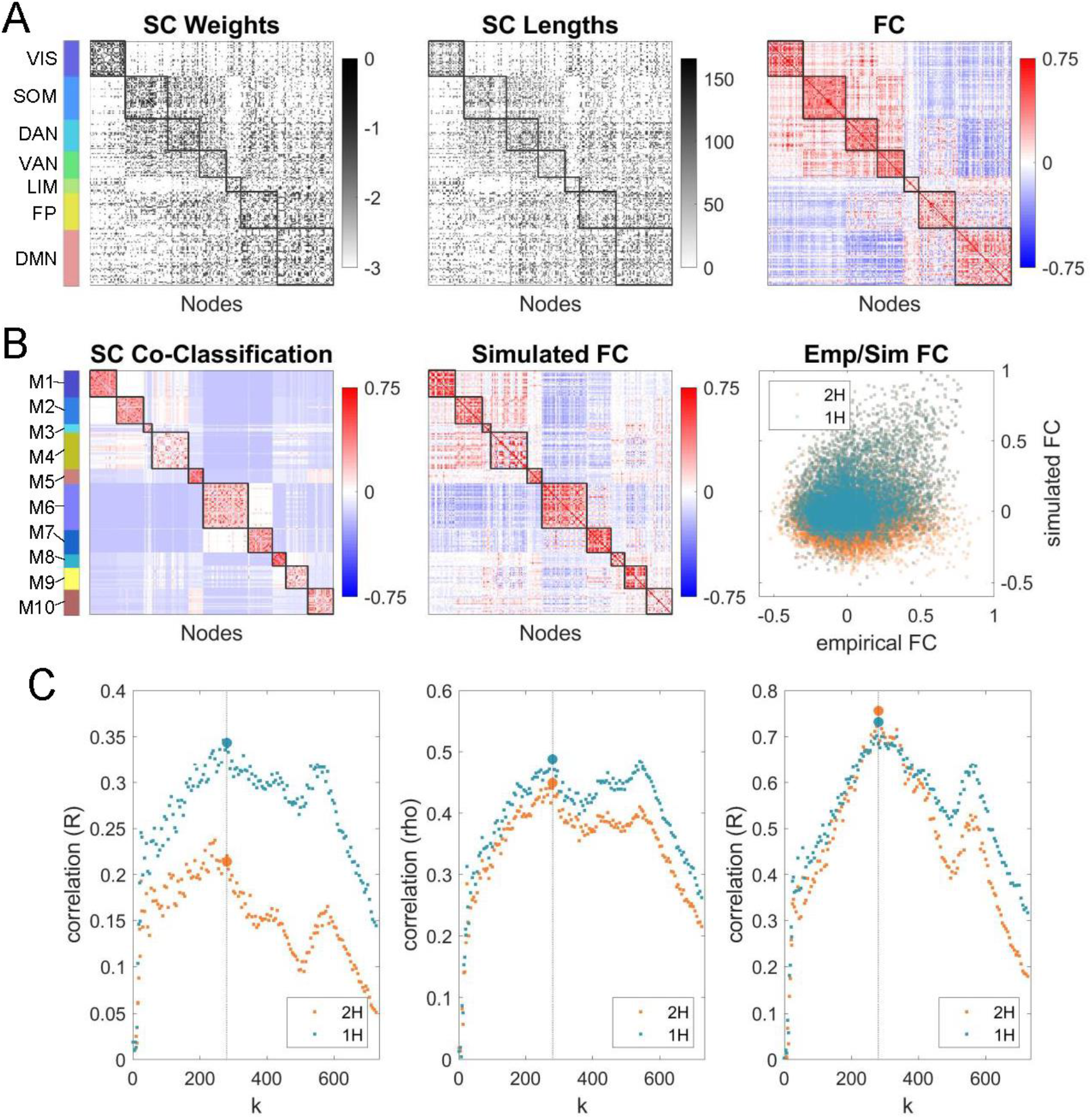
Structural and functional connectivity. (A) Empirical data. (left) SC consensus weight matrix; (middle) SC connection lengths (conduction delays); (right) FC, average of 95 subjects, 4 runs each; all panels shown in FC module node order (FC modules marked at left, cf. SI Fig 1A). (B) SC consensus and simulated FC. (left) Co-classification (agreement) matrix derived from the consensus SC matrix; (right) Simulated FC, average of 12 runs, *k*=280; both panels shown in SC consensus module node order (SC consensus modules marked at left, cf. SI Fig 1B). (right) Scatter plot showing comparison of empirical and simulated FC (orange: all node pairs; blue: intra-hemispheric node pairs only). (C) (left) similarity (Pearson correlation) between empirical and simulated FC, across all values of *k*; (middle) Spearman’s rho between empirical SC weights (*K*_*ij*_) and simulated FC; (right) similarity (Pearson correlation) between empirical SC co-classification matrix and simulated FC. All panels in (C) show data for the full range of the coupling parameter *k*, with orange dots indicating full-brain coverage (both cerebral hemispheres and their interconnections) and blue dots indicating intra-hemispheric connections only. Large dots indicate data for *k*=280 (stippled vertical lines), averaged over 12 runs.

Examining the RSS of simulated edge time series reveals brief, intermittent, high-amplitude peaks, or events, over a wide range of the coupling parameter *k* (Fig 3). In empirical data, events exhibit several characteristic features, including their disproportionate contributions to long-time estimates of FC and somewhat stereotypic topography [16]. Simulations reproduced these features (Fig 4). We tested whether the simulated edge time series exhibited some of the same features attributed to empirical edge time series. First, we constructed FC separately using either high- or low-RSS frames. We found that FC components constructed from high-RSS frames exhibited much greater similarity to the full FC estimate than those obtained from low-RSS frames, and high-RSS FC components were significantly more modular (Fig 4A,B). Additionally, the co-fluctuation patterns expressed during high-RSS frames were more similar to each other than low-RSS patterns, and e frames exhibited significant stereotypy across distant time points (Fig 4C,D).

**Figure 3:**
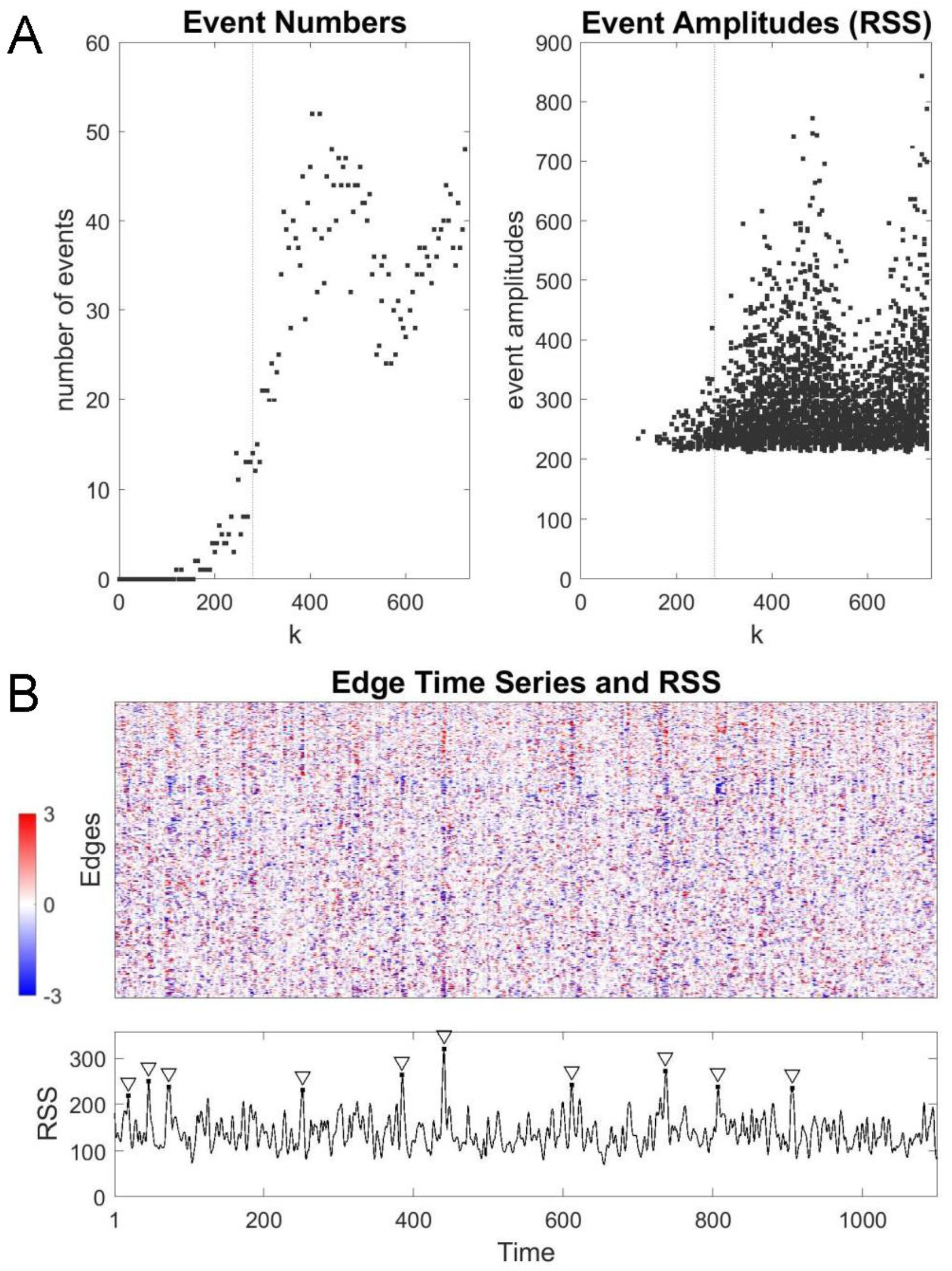
Events in simulated edge time series. (A) Number of events (left) and event amplitudes (RSS; right), over the full range of *k*. Application of the null model for event detection suppresses low RSS-amplitude peaks. (B) Example of simulated edge time series (*k*=280) and RSS amplitudes, with event peaks surviving null model comparison indicated by inverted triangles.

**Figure 4:**
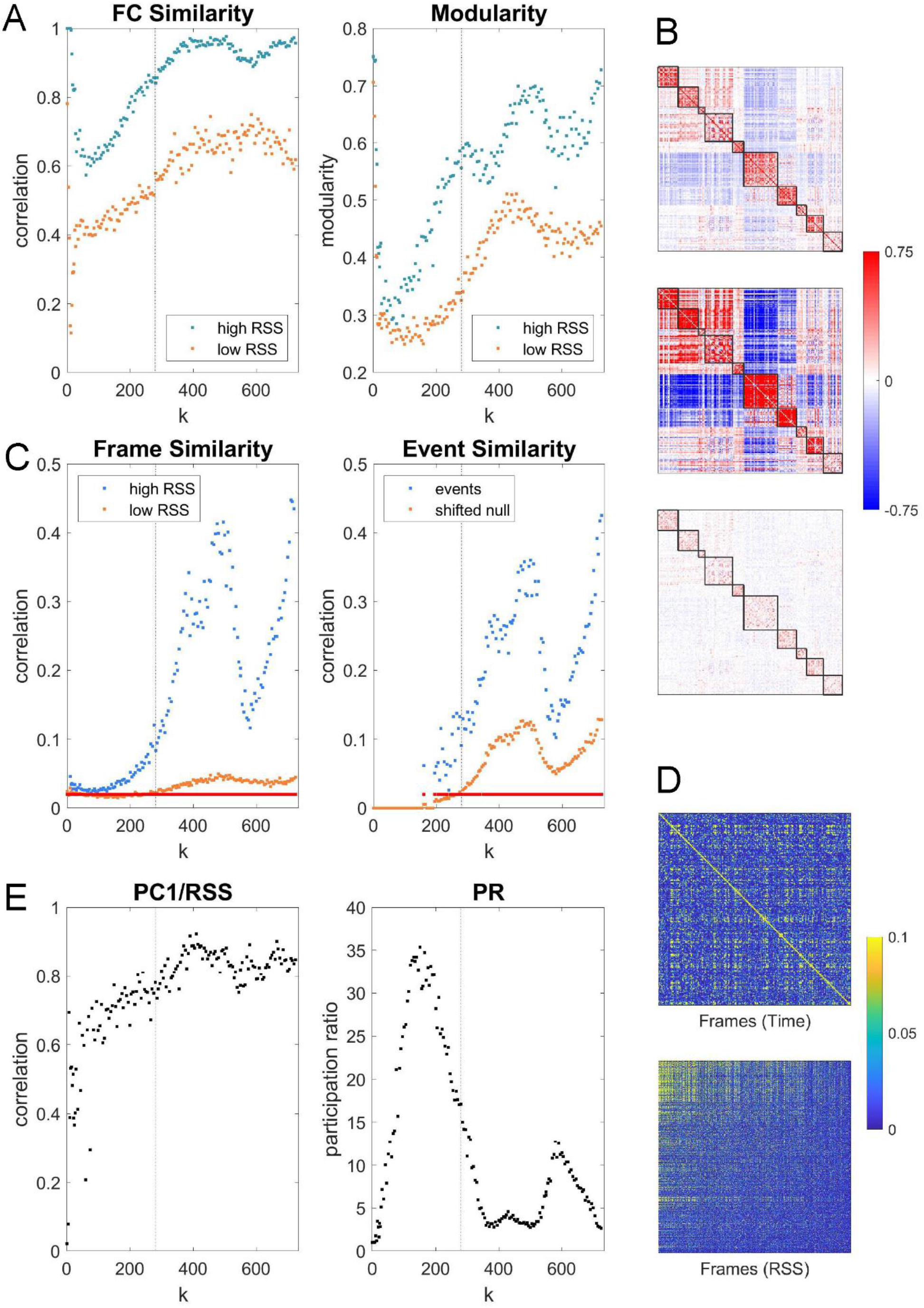
Properties of simulated edge time series. (A) Comparison of FC components derived from the top 10% (“high RSS”, blue dots) or bottom 10% (“low RSS”, orange dots) RSS frames. (left) Similarity (Pearson correlation) with full FC; (right) modularity. Both panels show data across the full range of *k*. (B) Example FC matrix (all frames, top; cf. Fig 2), and FC components (high RSS frames, middle; low RSS frames, bottom). Data averaged over 12 simulation runs of KS model, at *k*=280. (C) Similarity of frame sets sampled during high/low RSS epochs and during events. (left) Mean similarity (Pearson correlation) of frames within high/low RSS sets (110 frames each). Red dots indicate values of *k* for which significant differences between distributions were detected (Wilcoxon rank test, one-sided, *p* <0.001, uncorrected). (right) Mean similarity (Pearson correlation) of event frames compared to mean of 250 randomly offset frame sets. Red dots indicate values of *k* with *p* <0.001 (uncorrected). (D) Example plot of similarity of edge time series (Pearson correlation) across all frames within one simulation run (KS model, *k*=280; top: frames in original time sequence; bottom: frames sorted by RSS amplitude). (E) (left) Correlation of the largest principal component (PC1) of the edge time series with the RSS amplitude, across all values of *k*. (right) Participation ratio (dimensionality) computed from the FC covariance matrix. Stippled vertical line marks *k*=280 in panels A, C, and E.

We performed a principal components analysis (PCA) of edge time series, which yielded a distribution of principal components that differed in spatial pattern and temporal expression. The score (level of expression across time) of the largest principal component (PC1) was significantly correlated with global RSS amplitude (Fig 4E), suggesting that the PC1 captured a consistent component associated with high-RSS time steps. Consistent with the emergence of a significant PC1 component, the structure of the FC covariance matrix indicated the presence of low-dimensional dynamics (low participation ratio, *PR*) over most of the parameter range, coinciding with the occurrence of high numbers of events.

Next, we assessed the relationship of simulated events with the underlying structural connectivity. Data from 12 simulation runs (*k*=280) were aggregated to explore the topography of events and their temporal recurrence in greater detail (Fig 5). A total of 161 events, each represented by a vector of 19,900 edge co-fluctuation values, were extracted, and their mutual similarity (Pearson correlation) matrix was clustered to detect distinct sets of event patterns (Fig 5A). We tested whether patterns for the four largest event clusters were significantly aligned with SC consensus module boundaries (Fig 5B). The test employed a null model that rotates the structural modules on the cortical surface, thus approximately preserving their spatial contiguity (spin test, 50,000 rotations, ref [40]). Total co-fluctuation magnitude within all SC modules exceeded that obtained for all 50,000 null model rotations and for all four event clusters (*p* <0.00002). Testing for contributions of individual SC modules confirmed significant co-alignment of specific modules with co-fluctuation patterns (event cluster 1: SC modules M2, M4, M6; event cluster 2: M2, M6, M9; event cluster 3: M1, M2, M4, M6, M7; event cluster 4: M1, M4, M7; all *p* <0.0013, Bonferroni corrected). Different classes of events aligned with different subsets of structural modules and exhibited different time courses of co-fluctuations (Fig 5C). Our findings suggest that events belonging to different clusters are shaped by different combinations of modules present in the underlying structural connectivity.

**Figure 5:**
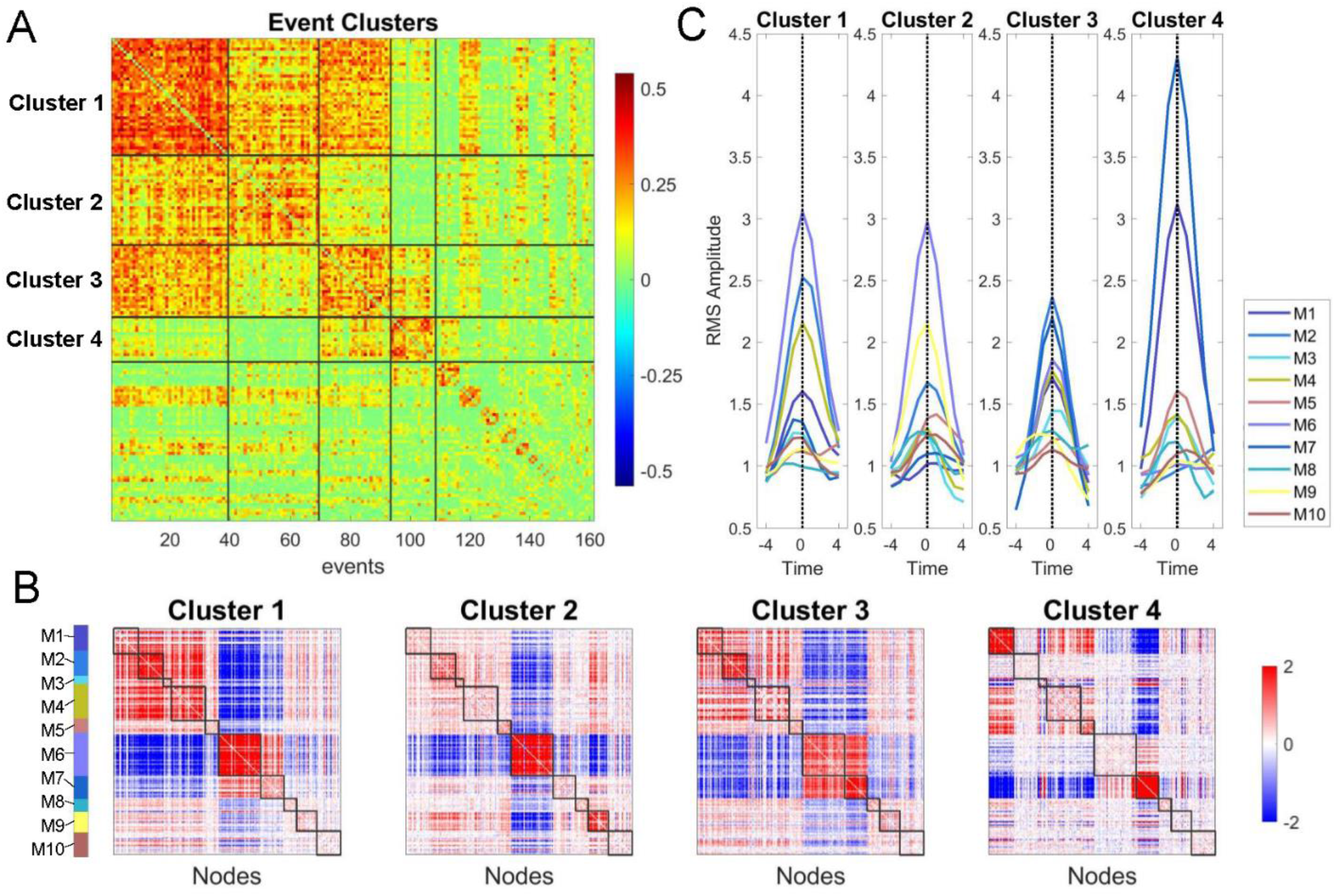
Event clusters and relation to SC consensus modules. (A) Clustered correlation matrix of event patterns (k=280, 161 events). Matrix is reordered to show event clusters, from largest to smallest. The top four clusters are delineated and contain 37, 30, 26, and 15 events, respectively. (B) Means of the events clusters (cluster centroids) displayed in matrix form, with nodes arranged in SC consensus order (cf. Fig 2). SC consensus modules for which the mean co-fluctuation significantly exceeded those obtained in a null distribution (spin test, 50,000 spins): M2, M4, M6 (cluster 1); M2, M6, M9 (cluster 2); M1, M2, M4, M6, M7 (cluster 3); M1, M4, M7 (cluster 4); all p<0.0013, Bonferroni corrected). (C) Mean time courses of RMS, computed for each SC consensus module, with time courses aligned to the event peak, for each of the four main event clusters (means of 37, 30, 26, 15 events, respectively). Time courses show mean co-fluctuation amplitude for each SC consensus module.

Several additional analyses were carried out to establish the robustness of these findings. Simulations employing different settings of the conduction velocity were analyzed to examine the frequency of events as well as the match between empirical and simulated FC (SI Fig 2). Findings indicated that events occurred over a wide range of velocities (6 m/s to 21 m/s), covering a range of plausible conduction velocities for inter-regional projections in primate cortex [41]. Events did not occur when simulations were carried out using a randomized SC matrix with near-absent modular organization (SI Fig 3). In contrast, events were readily observed in simulations that employed a synthetic (manually configured) SC matrix with strongly modular organization spanning specific node sets, and these events were aligned to the provided SC modular architecture (SI Fig 4). Finally, simulated BOLD time courses were generated by implementing a neural mass model (NMM, as employed in [28]) using the same SC consensus matrix and processed identically as for the KS model described above (SI Fig 5). The model matched empirical FC, exhibited robust structure-function correlations and showed events over a range of coupling parameters. NMM model events exhibited characteristics very similar to those found in empirical as well as KS model data. Importantly, as for the KS model, NMM event clusters were significantly aligned with structural consensus modules (SI Fig 5) (event cluster 1: M1, M4, M7; event cluster 2: M2, M4, M6; event cluster 3: M1, M2, M4, M6, M7, M9; event cluster 4: M1, M2, M3, M6, M7; all *p* <0.0013, Bonferroni corrected).

## Discussion

Here, we show that computational models of large-scale brain dynamics exhibit brief, intermittent, high-amplitude bursts of edge co-fluctuations (events) across broad ranges of coupling and conduction velocity parameters. Simulated events exhibit several characteristics that match those observed in empirical data, including strong similarity to full-length FC and recurrence across time. We find that simulated events display brain-wide patterns of co-fluctuation that are significantly aligned with network communities (consensus modules) present in the underlying structural connectivity. This relationship is reproduced in simulations that employ synthetic SC with defined modular architecture, is absent in randomized SC where modular structure has been degraded and is replicated in an independent model implementation. Overall, our study suggests a significant role for SC network modules in driving and shaping brief burst-like events that occur in fine-scale FC dynamics.

Recent studies have demonstrated that co-fluctuations of spontaneous brain activity as measured with fMRI exhibit brief, intermittent, and high-amplitude events [16–19]. These patterns contribute disproportionately to time-averaged FC, contain participant-specific information, and amplify brain-behavior correlations. However, their origins are poorly understood. We address this question using computational models, which have a long-established history as generative models of resting-state FC and FC dynamics [31]. Prior work has shown that their ability to match empirical data depends on the topology and weights of the underlying coupling matrix [28], noisy or chaotic dynamics and time delays [30], giving rise to variable, metastable dynamics, forming a rich repertoire of functional patterns. Here, we build on this body of work by extending model performance and analysis to fine temporal scales in BOLD time series. We show that model dynamic matches several recently described characteristics of empirically observed BOLD dynamics, including the recurrence of high-amplitude events in edge time series that drive full FC. Importantly, analyzing simulated functional connectivity with respect to the structural connectivity shows that specific patterns recurring in high-amplitude events are aligned with modular boundaries in the underlying SC matrix, suggesting a mechanistic basis of events in the topology of structural modules. This finding, in conjunction with the fact that events drive average long-time FC may account for the observation that modules form a consistent ‘core structure’ in functional connectivity [42]. Previous work has implicated ‘cluster synchronization’ in patterning of FC [32] and in temporal fluctuations related to near-critical and metastable dynamics [43]. A recent model [44] adopted the FC decomposition approach to track fine-scale co-fluctuation patterns linking them to neuronal cascades and nodal network structure. The model demonstrated that node centrality highly influences the node’s likelihood to participate in coordinated activations that may, in turn, spread within structurally densely connected clusters or communities. These findings are compatible with the role, proposed in the present study, of structural modules in shaping spatially organized co-activity and burst-like events.

The relationship between modular network topology and synchrony has been extensively investigated in the complex systems and networks literature [45–50]. In agreement with previous simulation studies, our model produced metastable dynamics and partial synchronization, which has been shown to manifest within clusters contained in the connectivity matrix. Applied to brain models, previous work demonstrated that slow fluctuations in modular FC topology on slower time scales (tens of seconds) depended on the integrity of structural modules [28] and that modular metastable states were driven by “cluster synchronization” [33]. Our findings suggest that these structural factors can act on fast time scales yielding burst-like events that carry signatures of structural connectivity. At intermediate coupling, networks with a strong community structure are conducive to fast, local synchronization, but not global synchronization [48]. Additionally, when moving toward full synchronization, smaller, more highly connected communities synchronize before larger communities, creating a mechanism by which network structure may impose different time scales on the system [51,52]. More generally, the finding that events are related to synchronization along the underlying modular structure resonates with this larger body of literature and supports the notion that the brain’s modular network architecture facilitates communication within modules while preventing global synchronization.

What causes events to occur? In simulations, we can confidently exclude scanner artifacts, spurious physiological signals, head movement, or variations in internal state as drivers of events. Indeed, analysis of empirical data has not found any significant correlation of timings or frequencies of events with nuisance variables related to acquisition, motion, or physiology [16]. Notably, in both KS and NMM models, the origin of events does not require a specific ‘built-in’ driving mechanism that triggers events at specific times. This observation is reminiscent of the emergence of complex dynamics on multiple time scales in high-dimensional neuronal systems [53]. While no forcing mechanisms are needed for events to occur, they are shown to depend critically on the topology of the SC matrix. We demonstrate that events are enabled by the presence of modular structural connectivity, and that the modular topology is imprinted in the specific dynamic patterns that manifest as events. We also emphasize that the model as presented here is implemented as a fully deterministic high-dimensional system of coupled differential equations that capture how elementary (microscale) processes give rise to emergent and collective (macroscale) system behavior. While many stochastic processes necessarily contain extremal events, our model is generative in a mechanistic, rather than purely stochastic, sense: observed fluctuations are localized in space and time, and their origin is rooted in the structure of the underlying anatomical network.

Our focus has been on extremal events, sharp excursions of global co-fluctuation amplitudes that drive FC. The cognitive relevance of such events is a current subject of investigation, with studies suggesting that fMRI event profiles carry subject-specific information [19]. Intriguing links may emerge between fMRI events and transient phase-locking of BOLD signals [54] and activity pulses [55], stereotypic patterns of propagation of intrinsic activity [56], and brain-wide traveling waves [57]. If events do support adaptive function, then we may speculate that specific network topological features have been selected to facilitate events. More work is needed to determine whether events have adaptive roles in promoting specific neuronal functions. For example, the brief and system-wide nature of events suggests relations to episodic synchronous neuronal activity such as avalanches [58] or ripples [59,60] that involve large multi-regional neuronal populations and may facilitate distribution of packets of neural information. The generative model furnished in this article may prove useful for examining and testing these and other hypotheses.

Limitations of the study should be discussed. Both model implementations documented in this article represent major simplifications of real anatomy and physiology. The models do not include subcortical regions and are carried out on connectivity data that is processed into a specific nodal parcellation at intermediate spatial resolution (200 nodes). Finer spatial scales of connectivity, more realistic dynamics, inclusion of region-specific model parameters [61], the addition of neuromodulatory systems [62], individualized connectomes, and geometric embedding of the underlying neural time series may improve model performance and match with empirical data. These limitations can be addressed in more accurate and data-driven multi-scale implementations, e.g., linking microscale anatomy and physiology to large-scale population responses.

Additional directions for future work, beyond increasing model realism, include application to task-driven dynamics, extrinsic perturbations, and the analysis of individual differences. Further understanding of the possible structural origins of events in empirical data may come from comparing empirical event patterns to SC modules. Because models give access not only to observed BOLD patterns but also to the underlying (phase-oscillator or neural population) time series sampled at millisecond resolution, fine-scale BOLD dynamics could be traced to fast fluctuations in synchrony at the microscale. These directions may provide additional valuable insights into the network basis of neuron-level dynamics that drive and shape functional connectivity.

## Acknowledgements

This material is based upon work supported by the National Science Foundation under Grant No. (NSF IIS-2023985; RFB, OS), National Science Foundation grant 1735095, Interdisciplinary Training in Complex Networks (MP) and Indiana University Office of the Vice President for Research Emerging Area of Research Initiative, Learning: Brains, Machines and Children (RFB). Data were provided, in part, by the Human Connectome Project, WU-Minn Consortium (principal investigators: D. Van Essen and K. Ugurbil; 1U54MH091657), funded by the 16 National Institutes of Health (NIH) institutes and centers that support the NIH Blueprint for Neuroscience Research and by the McDonnell Center for Systems Neuroscience at Washington University.

## Author Contributions

MP, MF, RFB and OS conceptualized and designed the experiments. MP and OS performed simulations and data analysis. MP, MF, RFB and OS made figures, wrote and edited the paper.

## Methods

### Dataset and Acquisition

All empirical data used for this study were derived from the set of 100 unrelated subjects acquired by the Human Connectome Project (HCP) [63]. Informed consent was obtained from all participants and all study protocols and procedures were approved by the Washington University Institutional Review Board. A detailed description of HCP data acquisition protocols can be found in [63,64]. All data were collected on a Siemens 3T Connectom Skyra equipped with a 32-channel head coil. Structural connectivity was derived from diffusion imaging and tractography. Briefly, subjects underwent two diffusion MRI scans, which were acquired with a spin-echo planar imaging sequence (TR = 5520 ms, TE = 89.5 ms, flip angle = 78°, 1.25 mm isotropic voxel resolution, b-values = 1000, 2000, 3000 s/mm^2^, 90 diffusion weighed volumes for each shell, 18 b = 0 volumes). These two scans were taken with opposite phase encoding directions and averaged. Functional connectivity was derived from resting-state functional magnetic resonance imaging (rs-fMRI), acquired with a gradient-echo echo-planar imaging (EPI) sequence over 4 scans on two separate days (scan duration 14:33 min; eyes open). Main acquisition parameters were TR=720ms, TE=33.1ms, flip angle of 52°, 2mm isotropic voxel resolution and a multiband factor of 8. Both SC and FC were mapped to regions (nodes) using a common parcellation scheme (N = 200 nodes in cerebral cortex; [38]), mapped to canonical resting state networks derived from [9].

For inclusion in the present study, we selected a subset of 95 (out of 100 total) subjects. Subjects were considered for exclusion based on the mean and mean absolute deviation of the relative root-mean square motion across either four resting state MRI scans or one diffusion MRI scan, resulting in four summary motion measures. If a subject exceeded 1.5 times the interquartile range (in the adverse direction) of the measurement distribution in 2 or more of these measures, the subject was excluded. These exclusion criteria were established before the current study. Four subjects were excluded based on these criteria. One subject was excluded for software error during diffusion MRI processing. The remaining subset of 95 subjects had the following demographic characteristics: 56% female, mean age = 29.29 +/- 3.66, age range = 22-36.

### Structural Preprocessing

Diffusion images were minimally preprocessed according to the description provided in [64]. Briefly, these data were normalized to the mean b0 image, corrected for EPI, eddy current, and gradient nonlinearity distortions, corrected for motion, and aligned to the subject anatomical space using a boundary-based registration [65]. In addition to this minimal preprocessing, images were corrected for intensity non-uniformity with N4BiasFieldCorrection [66]. FSL’s dtifit was used to obtain scalar maps of fractional anisotropy, mean diffusivity, and mean kurtosis. The Dipy toolbox (version 1.1) [67] was used to fit a multi-shell multi-tissue constrained spherical deconvolution [68] to the diffusion data with a spherical harmonics order of 8, using tissue maps estimated with FSL’s fast [69]. Tractography was performed using Dipy’s Local Tracking module. Multiple instances of probabilistic tractography were run per subject [70], varying the step size and maximum turning angle of the algorithm. Tractography was run at step sizes of 0.25 mm, 0.4 mm, 0.5 mm, 0.6 mm, and 0.75 with the maximum turning angle set to 20°. Additionally, tractography was run at maximum turning angles of 10°, 16°, 24°, and 30° with the step size set to 0.5 mm. For each instance of tractography, streamlines were randomly seeded three times within each voxel of a white matter mask, retained if longer than 10 mm and with valid endpoints, following Dipy’s implementation of anatomically constrained tractography [71], and errant streamlines were filtered based on the cluster confidence index [72].

The number of streamlines between nodes of the volumetric parcellations was recorded for each tractography instance. Fractional anisotropy, mean diffusivity, and mean kurtosis maps were sampled from the middle 80% of each streamline’s path, which were averaged within streamline and then across all streamlines between each pair of nodes. Streamline counts were normalized by dividing the count between nodes by the geometric average volume of the nodes. Since tractography was run nine times per subject, edge values had to be collapsed across runs. To do this, the weighted mean was taken, with weights based on the proportion of total streamlines at that edge. This operation biases edge weights towards larger values, which reflect tractography instances better parameterized to estimate the geometry of each connection.

A single group-averaged structural connectivity matrix was constructed by forming a consensus average preserving the length distribution of fiber tracts as well as matching the global connection density to the mean over all individual subjects [39]. The resulting matrix (200 nodes, 6040 connections, 30.4 percent density, 72.4 mm mean connection length) was used as a coupling matrix for the simulations. In addition to these group-averages, some simulations used SC matrices estimated on data from single participants.

### Functional Preprocessing

Minimal preprocessing of rs-fMRI data included the following steps [64]: i) distortion, susceptibility, and motion correction; ii) registration to subjects’ respective T1-weighted data; iii) bias and intensity normalization; iv) projection onto the 32k_fs_LR mesh; v) alignment to common space with a multi-modal surface registration [73]. This preprocessing resulted in ICA+FIX time series in the CIFTI grayordinate coordinate system. Additional preprocessing steps included: vi) global signal regression; vii) detrending and band pass filtering (0.008-0.08 Hz) [74]. Following confound regression and filtering, the first and last 50 frames of the time series were discarded, resulting in a final scan length of 13.2 min (1100 frames).

For comparing simulated to empirical FC, a single group-averaged FC matrix was derived by computing the mean FC over all 95 subjects and all 4 scans. When reporting correlations of full FC against other metrics, raw FC values were first passed through the Fisher z-transform.

### SC Consensus Clusters

Human connectome SC displays modular organization. To detect network communities in our SC consensus matrix we employed multiresolution consensus clustering [75] which allowed us to capture communities across multiple scales. The algorithm for modularity maximization was based on the Louvain method, employed a spatial resolution parameter *γ*, and operated in three stages. First, a coarse sweep (1000 steps) of the resolution parameter established outer bounds that yielded between 2 and N communities. Second, a finer sweep (10,000 steps) over this range collected partitions across the full range. These partitions were aggregated into a co-classification matrix, followed by subtracting a null model (constant across all node pairs) corresponding to an expected level of co-classification based on number and size of modules [75]. Finally, this null-adjusted co-classification matrix was re-clustered under a variable consensus threshold τ [76]. Resulting consensus partitions for different values of τ were collected. The most frequently sampled consensus partition contained 10 modules and was selected for subsequent analysis (SI Fig 1).

### Kuramoto-Sakaguchi Model Implementation

We implemented a version of the model that generates fast oscillatory dynamics and incorporates a coupling structure consisting of weighted undirected connections and the corresponding time delays, encoded in a phase delay (frustration) matrix. The fundamental equation of this Kuramoto-Sakaguchi (KS) model (Fig 1) is given as

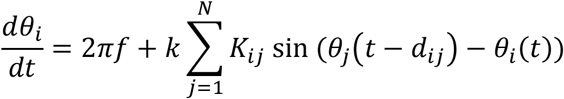

The global coupling parameter *k* was systematically varied. The mean natural frequency *f* was set to 40 Hz, with variability at each node (SD = 0.1 Hz). The coupling matrix *K*_*ij*_ corresponded, in most simulations, to the empirical SC consensus matrix (see above). A global scaling constant was applied to normalize *K*_*ij*_ for use in the simulations. The matrix of phase delays was derived from the physical lengths of each SC connection, computed as the group-averaged mean of its constituent streamlines, expressed in millimeters. Each SC connection length was converted to a time delay *d*_*ij*_ computed from a constant conduction velocity *ν*. Most simulations in the paper use ν = 12 *m/s*, resulting in a mean time delay of 6.1 *ms*, averaged over all connections in *K*_*ij*_. The conduction velocity was varied between ν = *m/s* and ν = 24 *m/s*, a range that was systematically explored in prior work [33] and roughly corresponds to the plausible physiological range for cortico-cortical projections in primates [41]. No noise was added to the phase time series.

The system of N coupled equations was integrated (using Matlab’s ode45 solver) at 1 ms time resolution, based on an implementation of the KS model in the Brain Dynamics Toolbox [77]. For each simulation, the initial condition consisted of phases drawn randomly and uniformly between [0, 2π]. Simulations proceeded for 812 s, which resulted, after removal of an initial 20 s transient, in 792 s of simulated time. Synchronization behavior of the system was tracked by computing the global order parameter

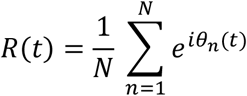

*R*(*t*) quantifies the global phase synchronization, with *R*(*t*) = 0 indicting complete asynchrony and *R*(*t*) = 1 indicating complete synchrony. After converting phases into signal amplitudes *θ*(*t*) (Fig 1C), the time series were convolved with a canonical haemodynamic response function (HRF; [77]; Fig 1D) resulting in synthetic BOLD time series (Fig 1E). These were lowpass filtered (0.25 Hz; [33]) and down-sampled at intervals of 720 ms, corresponding to the empirical TR in resting-state fMRI acquisitions, thus yielding a [*N* × *T*] simulated BOLD activity matrix, with T = 1100 time points (‘frames’). Finally, the global mean of these BOLD time series was regressed out, and the residuals were retained for computing edge time series and functional connectivity (Pearson correlation).

### Neural Mass Model Implementation

To demonstrate the robustness of our principal findings we configured a second dynamic model, closely based on earlier work [28]. Briefly, the neural mass model equations were adopted from a conductance-based model of neuronal dynamics designed to capture local population activity. Local populations of densely interconnected inhibitory and excitatory neurons whose behaviors are determined by voltage- and ligand-gated membrane channels were placed at each node, and the SC matrix provided inter-node couplings between excitatory neuronal populations. Parameter values were identical to those reported in [28]. No time delays were implemented, nor was stochastic noise added to the voltage time series. While other work has shown contributions made by time delays and noise to BOLD dynamics [30], here we were interested in whether a minimal set of ingredients (the SC weights and their modular network structure) could yield realistic event-like patterns. Only the global coupling parameter *k* was varied across runs.

Following numerical integration of the system of Nx3 coupled differential equations (using Matlab’s ode45 solver) and removal of a 20 s initial transient, the remaining time series were downsampled and converted to 792 s and 1100 frames of synthetic BOLD data, as described above for the KS model. All analyses were carried out identically for BOLD time series derived from both KS and NMM models.

### Edge Time Series and Co-Fluctuation Amplitude

Simulated BOLD time series were processed into edge time series as previously described for empirical fMRI data [16–19]. Nodal BOLD signals were z-scored, and edge time series (one for each node pair) were computed as the element-wise product of the respective node time series:

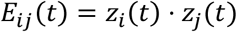

The mean of each edge time series is exactly equal to the corresponding Pearson correlation (functional connectivity), i.e., 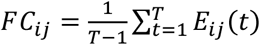. Edge time series, representing edge co-fluctuations, formed a [*K* × *T*] matrix, with *K* = 19900 (for *N* = 200) and *T* = 1100. The system-wide amplitude of all co-fluctuations was computed as the ‘root sum square’ (RSS) or, in cases where different-size node sets were compared, as the ‘root mean square’ (RMS).

Prior work in empirical fMRI data established that FC components derived from high RSS frames have higher similarity to full FC and have higher modularity than those derived from low RSS frames [16]. We computed FC components for subsets of frames corresponding to the top and bottom deciles of RSS (‘high’ and ‘low’ RSS), by taking the means across the respective subsamples of the edge time series. Modularity was computed on the resulting FC component matrix by applying modularity maximization (Louvain algorithm) adapted for use with matrices that contain both positive and negative edges [78]. The modularity metric *Q*^*^ summed contributions by positive and negative edges as

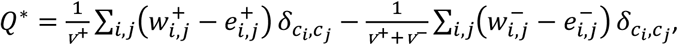

where 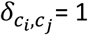 if nodes *i* and *j* are within the same community and 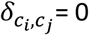 otherwise. The positive and negative superscripts to the edge weight *w*_*i,j*_ between nodes *i* and *j* are used for separating positive and negative edge weights, where 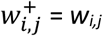 and 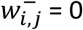 if *w*_*i,j*_ > 0, and 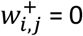 and 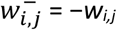 otherwise. The term 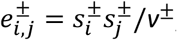, where 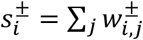 and 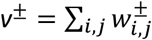, is the expected positive or negative edge weight between nodes *i* and *j* if edges were randomly distributed. All edges were retained, and the value of the modularity metric was computed while setting the resolution parameter to the default of *γ* = 1. Very similar results were obtained for other (higher and lower) settings of *γ*.

Another previous finding indicated that frames obtained when RSS is high were more similar (stereotypic) across time than low RSS frames [16,18,19]. In the simulation data, two related tests were performed to examine the similarity of high/low RSS frames as well as the similarity of event frames versus a time-shifted null. High/low RSS frame sets were extracted as described above and the distribution of all pairwise Pearson correlations among their constituent edge co-fluctuation patterns was compared (Wilcoxon rank sum test for equal medians, one-tailed, *p* <0.001, uncorrected). Event peaks were extracted as described below, and the set of all Pearson correlations among these events was compared against 250 sets computed from an equal number of randomly shifted time points, with offsets chosen uniformly from the interval [–1100 1100]. The mean correlation of the actual events was compared against these 250 random values and considered significant if *p* <0.01 (uncorrected; actual value exceeds null model at least 247/250 times).

Two additional analyses delivered insights on the relation of edge time series patterns and RSS, as well as on the dimensionality of the edge dynamics. First, performing principal component analysis (PCA) on edge time series yields a series of principal components (PCs). The correlation between the scores of the largest PC (PC1; taken to represent its expression over time) and the RSS is computed as Spearman’s ρ. Second, the eigen-decomposition of the covariance matrix of the neural activity (BOLD time series) yields a series of eigenvalues {λ_*i*_} from which the participation ratio (*PR*) can be computed as a measure of the dimensionality of the system [79]:

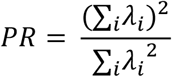

The value of *PR* is highly correlated with the cumulative explained variance of the largest principal components of a PCA decomposition of the BOLD covariance matrix.

### Event Detection and Statistics

Peaks in the RSS statistic that exceed a statistical criterion are termed events. To determine statistical significance, we performed non-parametric permutation tests on each model run ([19]; Fig 1G). BOLD time series were randomly shifted (using Matlab’s circshift operator), each separately with an offset chosen uniformly from the interval [–1100 1100], thus approximately preserving each node’s autocorrelation, while randomizing the cross-correlation between nodes. This null model was applied 1000 times to each model run, and edge time series and RSS were computed for each instance. Time points in the original time series for which the empirically observed RSS amplitude exceeded all RSS values for all null model instantiations (*p* <0.001) were retained. The intersection between all peaks in the original RSS and these time points constituted significant events in the model run. Event amplitudes corresponded to the RSS value at the time each such peak occurred.

At very low coupling (*k* ≤ 35 in the KS model), we observed occasional excursions in single-node time series which resulted in sharply fluctuating BOLD signal amplitudes and extremal z-scores. Extreme z-scores in single or very few nodes can result in sharp RSS spikes at the system level due to the transitive and multiplicative nature of the edge time series calculation. This phenomenon did not match characteristics of events observed in empirical fMRI data. Hence, RSS peaks that coincided with excursions of at least one BOLD time series above or below z = ±4.5 (the 99.99^th^ percentile of empirically observed BOLD signal amplitudes) were excluded from the analysis. Such peaks occurred infrequently (at most once per 1100 frames) at couplings *k* > 35.

### Event Clustering

Once statistically valid events were detected, their corresponding edge co-fluctuation patterns were extracted and aggregated across multiple runs (*k*=280 for the KS model; *k*=0.175 for the NMM model; 12 runs each). The Pearson correlation matrix for these event patterns was clustered using a version of modularity maximization (employing the Potts model) and multi-resolution consensus clustering [75]. The resolution parameter was stepped through a range of 10,000 values covering modular partitions yielding between 2 and N modules. All partitions were aggregated into a single co-classification matrix which was scaled between [0 1] followed by subtraction of the analytic null [75] and subjected to consensus clustering with τ = 0.

The four largest clusters were retained for subsequent analyses (labeled event clusters 1-4). A single mean edge co-fluctuation pattern (cluster centroid) was computed for each cluster. Applying the SC consensus module partition to the four centroids yields a representation of how event-related co-fluctuations distribute relative to the structural connectivity. To test whether mean co-fluctuations within each of the structural modules were significantly above or below chance, the modular components of edge co-fluctuations were recomputed using the spin test [40] which rotates the structural modular partition 50,000 times on the cortical surface. This procedure preserves not only the number but also (approximately) the spatial proximity of nodes contained in the original SC consensus modules. SC modules were considered aligned with event patterns if their internal co-fluctuations were greater than those from the 50,000 rotations, at a Bonferroni-corrected 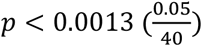.

### SC Randomization and Synthetic SC

The SC consensus matrix was randomized using a rewiring algorithm [80] that preserved the degree sequence exactly and the strength sequence approximately. Connection lengths (delays) were preserved at each connection such that the connection weight/delay relationship was retained. The resulting randomized matrix matched the SC consensus in all summary statistics pertaining to weight or delays, while comprising a globally randomized architecture. The modular architecture, as revealed through SC consensus clustering (SI Fig 3), was significantly muted.

A synthetic SC matrix was constructed such that the modular arrangement of the connections was predetermined (4 modules in the case of the matrix displayed in SI Fig 4) while retaining some summary statistics of the empirical SC consensus matrix (density). The matrix was used in KS model simulations with a rescaled coupling parameter of *k* = 25. This rescaling ensured that the global coupling strength provided by the synthetic SC matrix approximately matched that provided by the SC consensus at *k*=280.

## Supplementary Information

**SI Figure 1:**
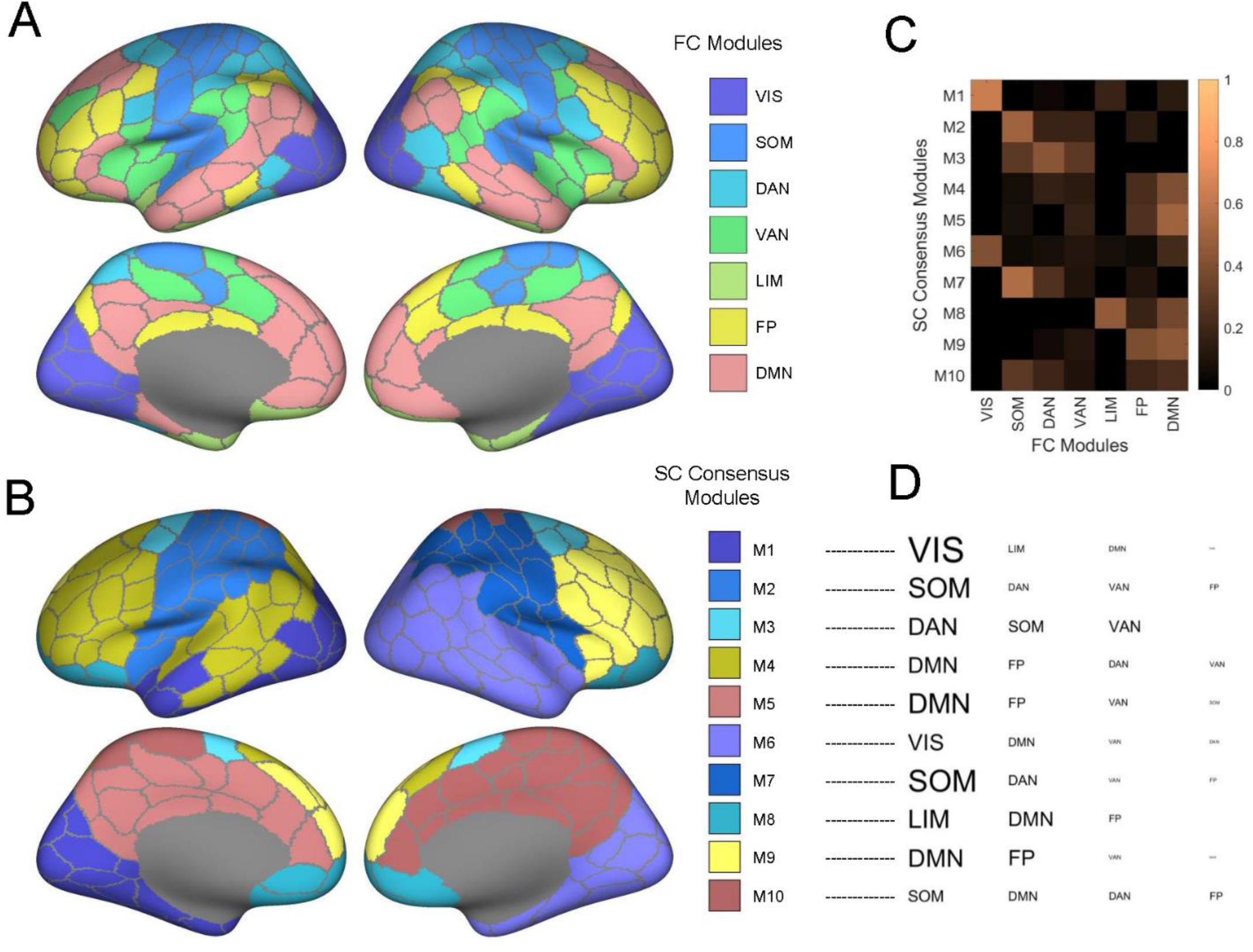
Parcellation and modular partitions. (A) Canonical functional systems (FC modules, see refs. [9,38]). (B) SC consensus modules: 10 modules derived from multi-resolution consensus clustering of the SC consensus matrix. SC consensus partition and Yeo partition are significantly related (mutual information between SC consensus and FC modules: MI = 0.277; mean/SD of 10000 randomly shuffled SC consensus partitions vs. FC modules: MI = 0.075 ± 0.013). (C) Comparison of FC modules and SC consensus modules. Matrix plots mutual overlap, with rows summing to 1. (D) FC modules that most closely match the SC consensus modules, based on the overlap matrix in panel (C). Font size indicates relative magnitude of overlap.

**SI Figure 2:**
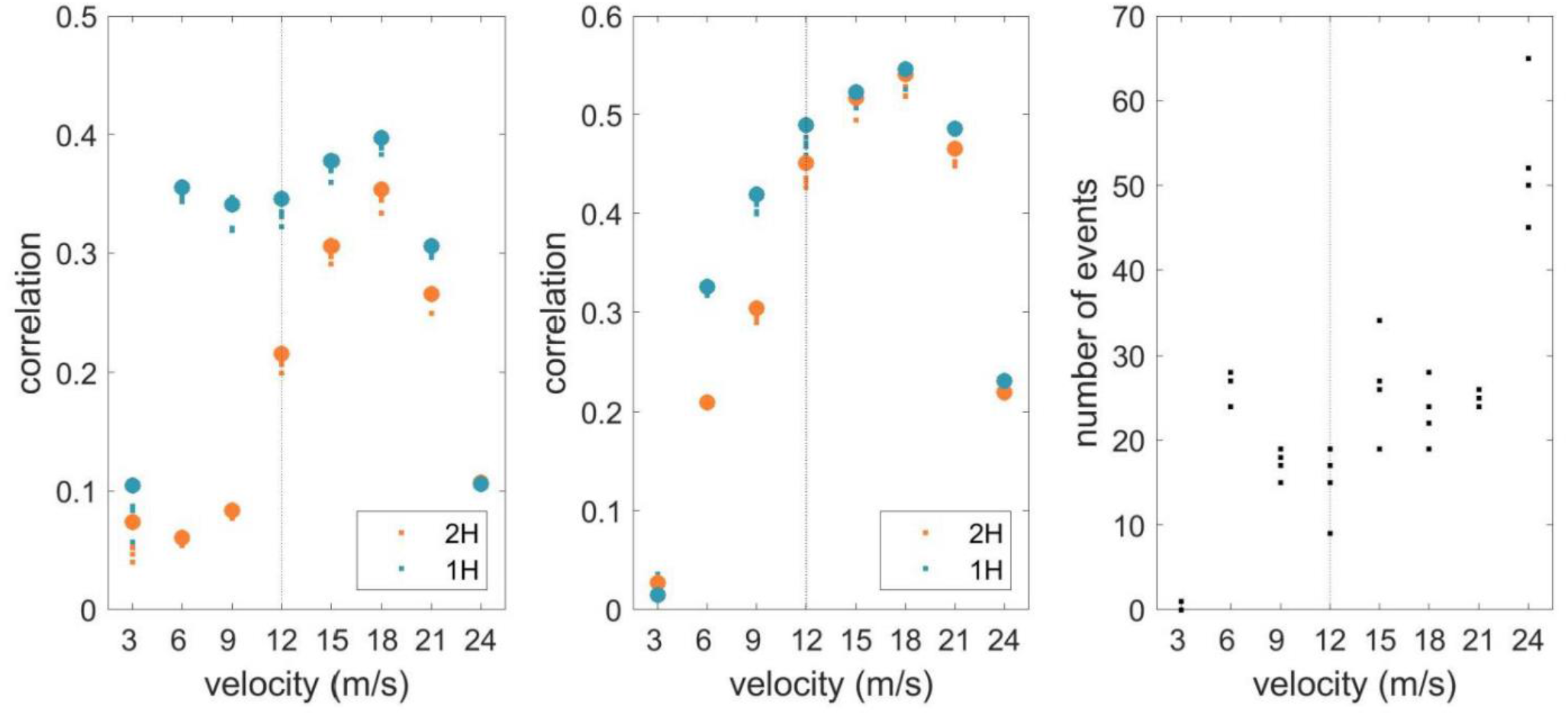
FC and events at different settings of the conduction velocity. All other simulation parameters are identical to the simulations reported in the main text. Data are shown for 4 simulation runs at each setting of the velocity. Large dots show means over all 4 runs. (left) Similarity (Pearson correlation) between empirical and simulated FC; (middle) Spearman’s ρbetween empirical SC weights (*K*_*ij*_) and simulated FC; (right) Number of events.

**SI Figure 3:**
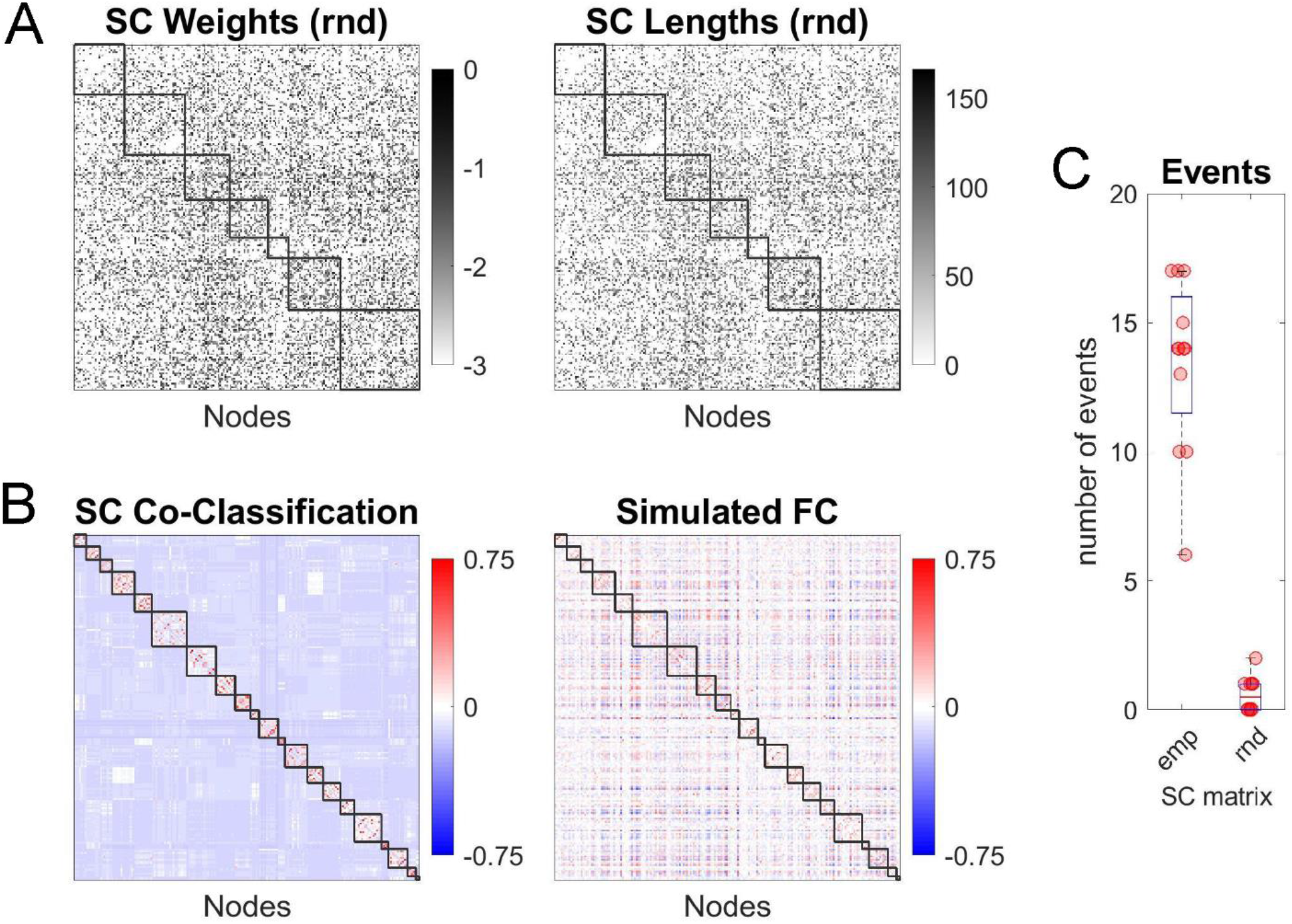
Randomized SC matrix. (A) SC weights (left) and connection lengths (right). Randomization preserves node degree and approximates node strength. Weight and length distributions are identical to those in the empirical SC matrix. (B) SC co-classification matrix (left) and simulated FC (right; 12 runs, *k*=280). (C) Event counts for 12 runs using the empirical SC (‘emp’) and the randomized SC (‘rnd’).

**SI Figure 4:**
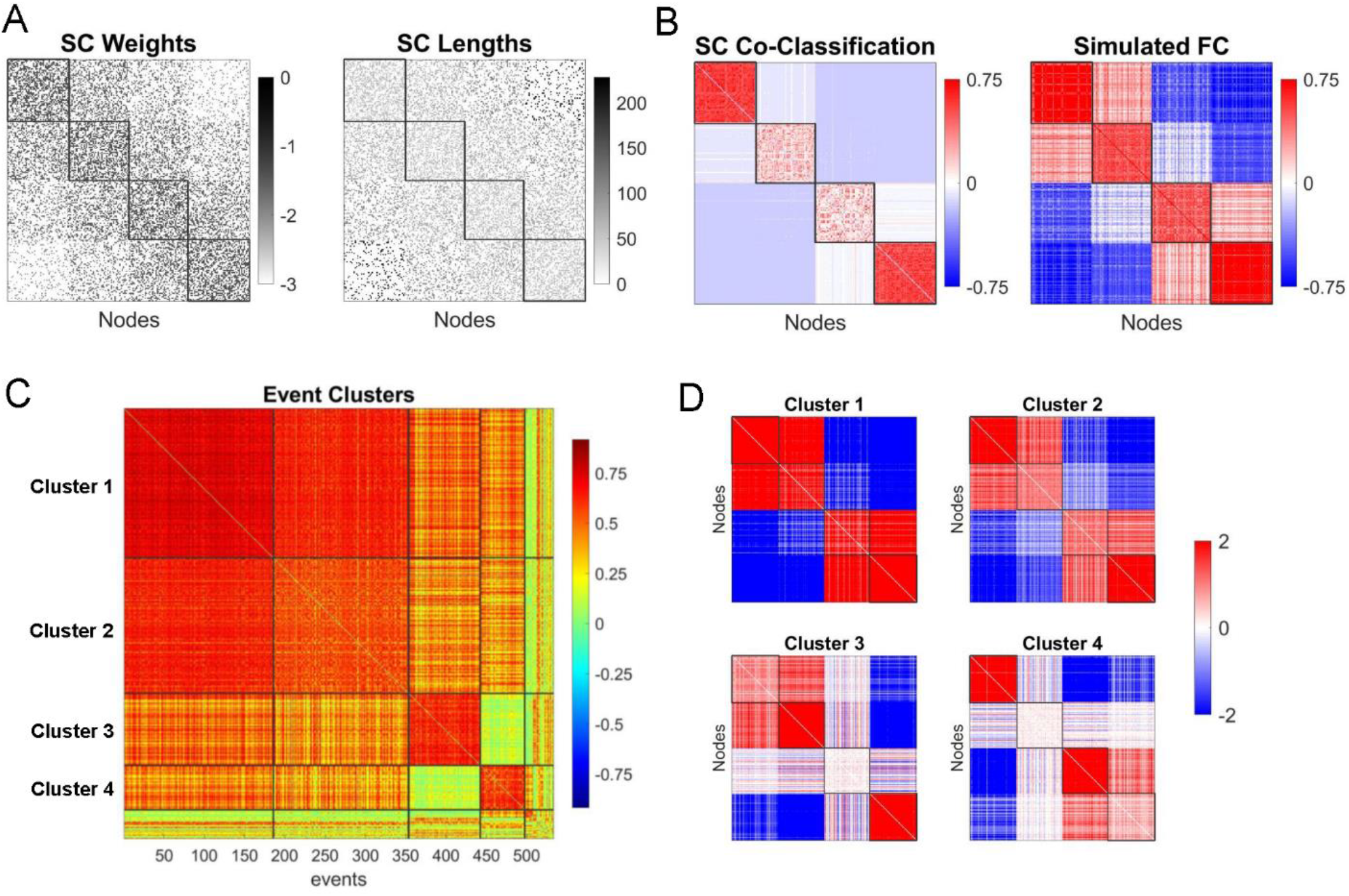
Synthetic matrix. (A) SC weights and lengths, arranged into 4 structural modules. (B) SC co-classification and simulated FC (mean of 12 runs, *k*=280). (C) Correlation matrix of event frames detected across 12 runs (*k*=280), with the four largest clusters indicated (185, 168, 89, 55 events, respectively; total: 533 events). (D) Edge co-fluctuation patterns for the four main event clusters (means shown, corresponding to cluster centroids). SC consensus modules for which the mean co-fluctuation significantly exceeded those obtained in a null distribution (spin test, 50,000 spins), numbered from the top of the matrix: M1, M4 (cluster 1,2); M2, M4 (cluster 3); M1, M3 (cluster 4); all *p* <0.0031, Bonferroni corrected).

**SI Figure 5:**
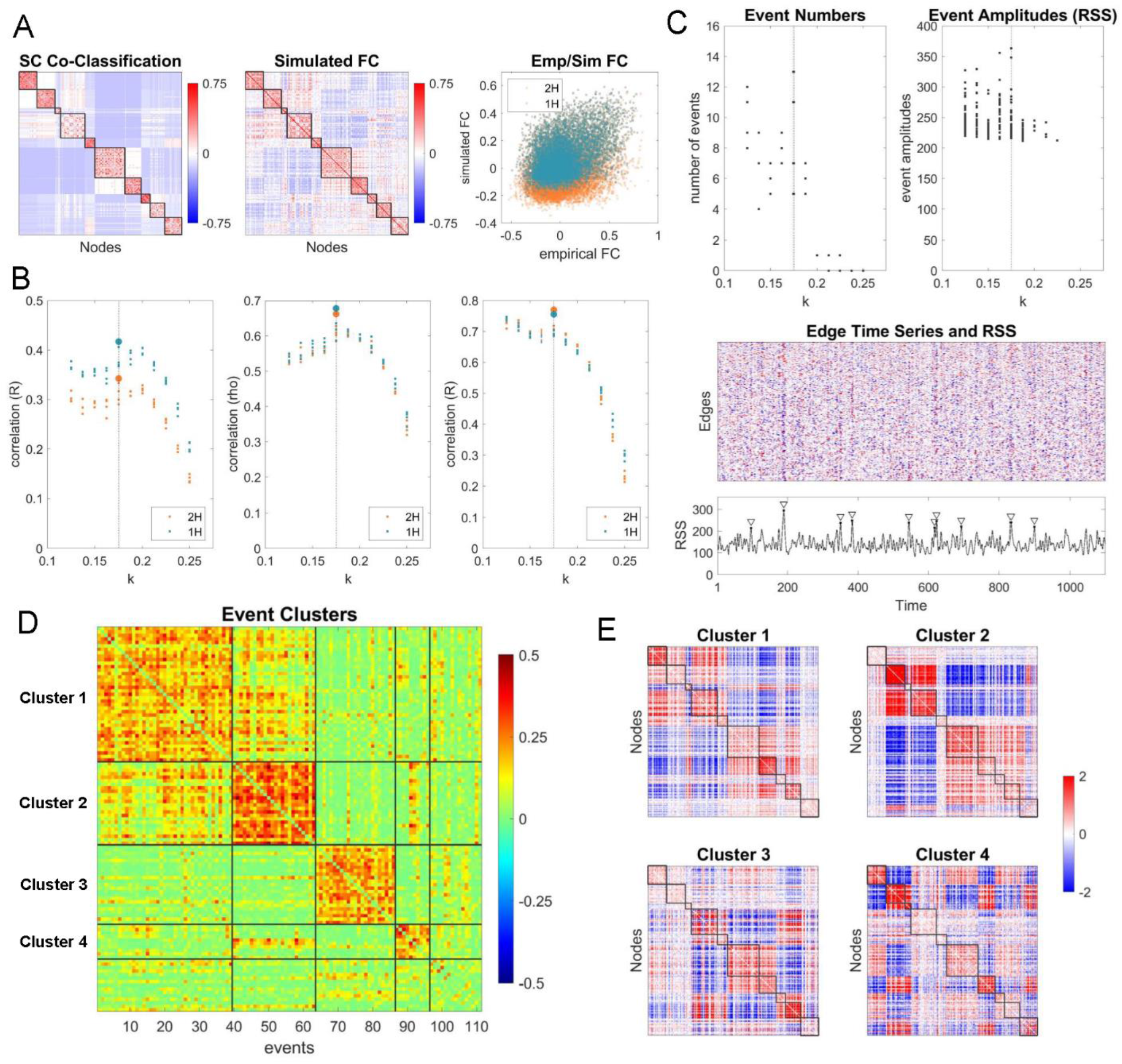
NMM simulations. (A) SC co-classification (right panel; cf. Fig 2) and simulated FC (middle panel; mean of 12 runs, *k*=0.175). Scatter plot (right panel) shows relation of empirical to simulated FC (mean of 12 runs, *k*=0.175), for all node pairs (orange dots; *r*_*a*_= 0.342, *p* =0) and inter-hemispheric node pairs (blue dots; *r*_*i*_ =0.417, *p* =0). (B) Pearson correlation between simulated and empirical FC (left panel; k=0.175 indicated by stippled line), Spearman correlation of simulated FC with empirical *K*_*ij*_ (middle panel), and Pearson correlation of simulated FC with SC co-classification (right panel). (C) Event numbers and amplitudes (top panels) over a range of *k*. Sample edge time series and RSS plot (*k*=0.175; bottom). Events indicated by inverted triangles. (D) Correlations of 111 events, with four main event clusters indicated, comprising 39, 24, 23 and 10 events, respectively. (E) Means of the events clusters (cluster centroids) displayed in matrix form, with nodes arranged in SC consensus order (cf. Fig 2, Fig 5). SC consensus modules for which the mean co-fluctuation significantly exceeded those obtained in a null distribution (spin test, 50,000 spins): M1, M4, M7 (cluster 1); M2, M4, M6 (cluster 2); M1, M2, M4, M6, M7, M9 (cluster 3); M1, M2, M3, M6, M7 (cluster 4); all *p* <0.0013, Bonferroni corrected).

